# E-HAPLOS: Electrical Impedance Human-guided Assessment with Pressure for Lump Observation System

**DOI:** 10.64898/2026.07.17.738661

**Authors:** Ralph Jirell E. Posio, Kirsten G. Magpili

## Abstract

Breast cancer is the leading cause of cancer-related deaths among women in the Philippines. Over 65% of these cases are diagnosed when they are advanced (Montemayor, 2023). This highlights the need for improved early screening devices. E-HAPLOS, or Electrical Impedance Human-guided Assessment with Pressure for Lump Observation System, is a low-cost glove with sensors designed to improve early detection of suspicious breast lump through touch. It integrates force-sensitive resistors (FSRs) to measure tissue stiffness and Electrical Impedance Spectroscopy (EIS) to analyze conductivity across different frequencies—properties that are closely linked to breast cancer.

The prototype uses an ESP32 microcontroller that transmits real-time pressure and impedance data to the website. Tested on gelatin breast models with simulated lump, the FSRs effectively identified lump locations by recording higher mean force values (45.81 kPa vs. 33.57 kPa). This guided approach allowed the combined FSR-EIS system to reach a diagnostic performance with an Area Under the Curve (AUC) above 0.94, a significant improvement over unguided measurement (AUC ≈ 0.78). A two-way ANOVA confirmed a significant difference in diagnostic performance based on the system modality (p < 0.001). Tukey’s Honesty Significant Difference (HSD) test showed that the FSR-EIS system was statistically superior to both the unguided EIS (p < 0.001) and FSR-only system (p = 0.041). Results demonstrate the synergistic effect of the integrated system, enabling accurate differentiation of suspicious lumps from normal tissue.

The FSR-EIS system of the E-HAPLOS glove shows a great potential for detection of lumps in simulated breasts as a screening tool.

## INTRODUCTION

According to the American Cancer Society (2024), breast cancer is the most diagnosed cancer among women. It occurs due to changes in breast tissue and the uncontrolled division of cells that typically results in a lump or mass. The majority of breast cancers begin in milk glands (lobules) or in the tubes (ducts) that connect milk glands to the nipple. The consequences of late diagnosis are severe, as 65% of cases are diagnosed in advanced stages (III or IV), leading to a five-year survival rate of just 44.4% Montemayor, (2023). This is in stark contrast to countries with robust early detection programs, highlighting a critical gap in accessible healthcare technology. The low coverage of breast cancer screening in the Philippines, reaching only about 1% of women, is primarily due to high costs, limited availability of advanced diagnostics in rural areas, and the fear of a difficult diagnosis Pazzibugan, (2023).

Early tumors can appear as stiffer lumps of tissue which can be detected through hand palpitation. They also exhibit distinct electrical conductivity characteristics, as the electrical properties of malignant tissues differ significantly from those of benign tissues. Electrical Impedance Spectroscopy (EIS) can precisely measure these differences by applying currents at various frequencies, revealing subtle variations imperceptible to touch. However, the subjectivity of manual palpation may lead examiners to overlook these subtle signs, especially those not experienced in examining for lumps Zhou et al. (2017); Shimatani et al. (2022).

By providing an affordable early screening method that combines the accessibility of physical examination with the objectivity of technology-based tools, this device bridges the gap between manual palpation and high-cost imaging. E-HAPLOS offers a comprehensive screening tool particularly valuable for resource-limited settings where traditional diagnostic methods are unavailable by exploiting suspicious breast lumps’ characteristics.

In addition, the study will align on the United Nation’s Sustainable Development Goals, Specifically Goal 3 (Good Health and Well-Being), promoting affordable and innovative medical solutions, Goal 7 (Affordable and Clean Energy) by creating a low-power and energy-efficient device for areas with limited electricity, and Goal 9 (Industry, Innovation and Infrastructure) by developing medical technology and contributing to accessible healthcare infrastructure.

E-HAPLOS: Electrical Impedance Human-guided Assessment with Pressure for Lump Observation System addresses this gap by integrating force-sensitive resistors (FSR) for stiffness measurement and Electrical Impedance Spectroscopy (EIS) for electrical property analysis, all into a single wearable glove. This multi-sensor approach transforms subjective touch into quantifiable, objective feedback through a website interface, enabling users to identify abnormal patterns with greater accuracy. The combined FSR-EIS system and individual sensor modalities in detecting simulated breast lumps.

## METHODS

### Research Design

This study employs an experimental research design to assess the synergistic effect of the E-HAPLOS glove, integrating EIS and FSR, in detecting suspicious breast lumps. Two groups were formed: a control group that examined a healthy breast model, and an experimental group that examined a model with a simulated palpable lump of a suspicious mass. Both groups underwent the same procedure under controlled conditions, with standardized materials, instructions, and time allocation. Data collection was conducted through observation checklists and scoring sheets, while analysis involved descriptive statistics, ROC–AUC evaluation, and logistic regression analysis to identify relationship of FSR and the lump and significant differences between the two groups. Two-way ANOVA using the AUC scores will be used to identify the significant difference between the groups then use Tuckey’s HSD to classify the group that made the significant difference.

**Figure 1.**
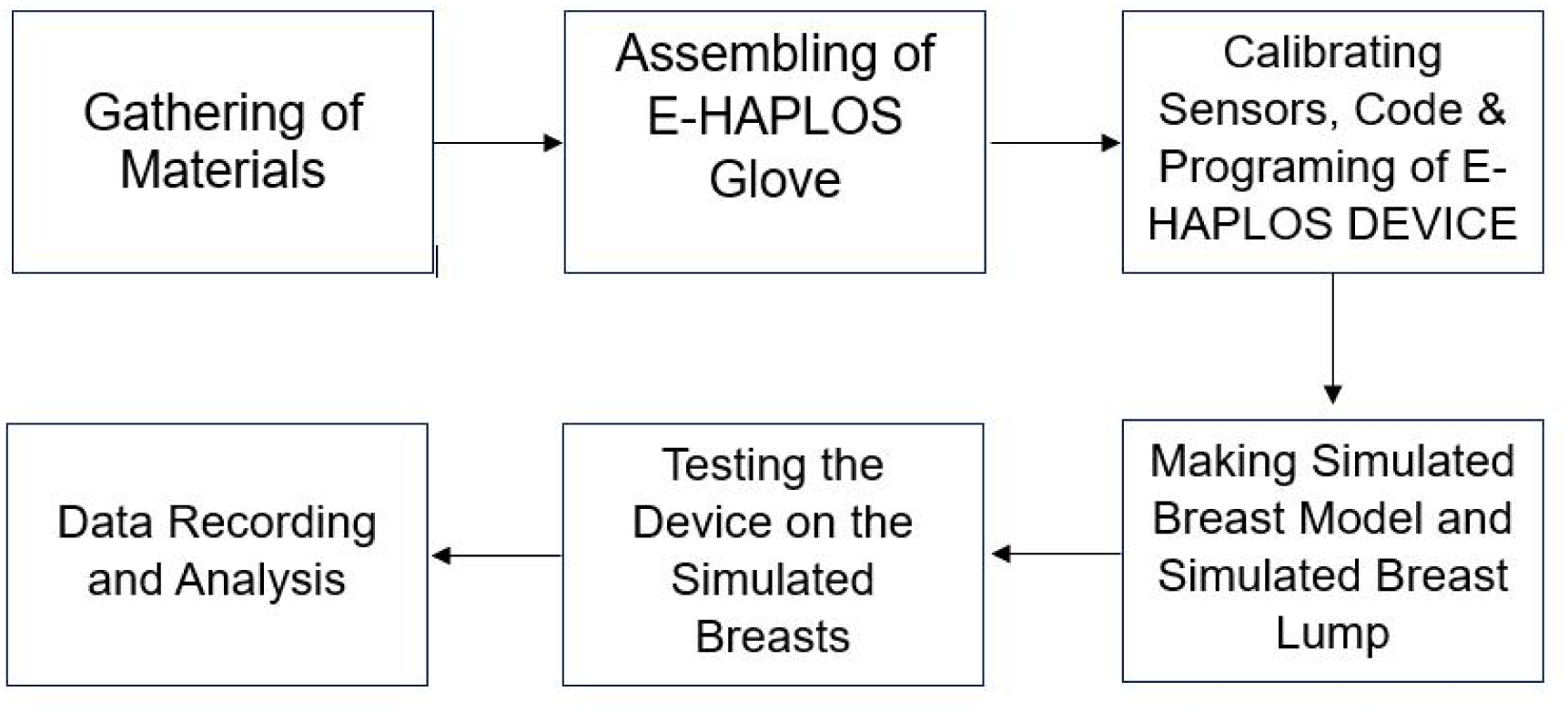
Methodological Process of E-HAPLOS

### I. Gathering of Materials

All necessary electric components and materials for the construction of the E-HAPLOS prototype were collected. These included an ESP32, microcontroller, an ADS1115 analog-to-digital converter, an OPA2143 operational amplifier, two force-sensing resistors (FSRs), two stainless press button fastener, and assorted jumper wires, stranded wires (22 AWG), solder, and a textile glove, additionally, materials for the simulated breast phantoms were prepared, including gelatin, agar, salt (NaCl), and graphite dust.

### II. Assembling of the E-HAPLOS Glove

The prototype was constructed on a washable, stretchable textile glove. The two FSRs were secured on the fingertips at the index and middle finger to measure localized pressure and stiffness. The two EIS electrodes were also placed on the fingertips, adjacent to the FSRs, to measure the electrical properties of the tissue. All components running through the glove were interconnected via thin stranded wires to ensure comfort and flexibility. To ensure stable operation and minimize high-frequency and frequency-dependent noise, extensive power supply decoupling is implemented. Multiple 100nF ceramic capacitors are placed across the VCC and GND pins of every active Integrated Circuit (IC): specifically, between the power (VDD) and ground (GND) pins of the ESP32, ADS1115 (Pin 4 VDD to Pin 3 GND), OPA2134 (Pin 8 V+ to Pin 4 V-), and HD14051 (Pin 16 VDD to Pin 8 VSS/GND). A 0.1nF ceramic capacitor is also connected from the middle wiper of the 2.5V potentiometer to ground, stabilizing this critical analog reference. All these circuits are enclosed in a cage lined with aluminum foil to provide EMI shielding, with the foil connected to system ground.

### III. Calibrating of Sensors, Code & Program of E-HAPLOS Glove

The E-HAPLOS system’s functionality relies on the precise interaction of its electronic components, governed by firmware, and the accurate calibration of its sensors. This section details the system’s operational principles, hardware implementation, sensor calibration procedures, and firmware logic.

The E-HAPLOS system integrates Electrical Impedance spectroscopy (EIS) for sensing electrical properties and Force-sensing Resistors (FSRs) for pressure distribution monitoring.

#### A. Electrical Impedance Spectroscopy (EIS)

EIS non-invasively sense the electrical properties of the simulated breast models. The process involves:

1. A small, precisely generated alternating current (AC), varying across a sweep of frequencies from 100kHz to 500kHz, is injected to the phantom via a pair of electodes (I+ and I-).
2. A separate, non-contacting pair of electrodes (V+ and V-) measure the resulting voltage across the phantom. This four-electrode configuration is important to minimize errors from electrode-skin contact impedance, ensuring that the measured voltage accurately reflects the tissue’s intrinsic electrical properties.
3. From the known injected current and the measured voltage, the system calculates the tissue’s electrical impedance (a complex quantity with magnitude and phase). By measuring this across various frequencies, the electrical properties of the phantom are obtained, which changes with physiological state.

##### a Transimpedance Amplifier (TIA) Circuit

The core of the EIS measurement chain is the Transimpedance Amplifier (TIA) circuit, built around an OPA2134 operational amplifier. One part of the OPA2134 acts as a voltage-controlled current source, converting the ESP32’s generated voltage signal into a precise AC current for injection. Another part of the OPA2134, configured with a feedback resistor (10 kΩ) and capacitor (30 pF), functions as the TIA, converting the minute current response from the tissue into a measurable voltage that the Analog-to-Digital Converter (ADC) can process. This circuit is crucial for sensitive and accurate current-to-voltage conversion.

##### b Force-sensing Resistor (FSR)

Two FSRs are integrated into the glove to monitor pressure. There electrical resistance changes proportionally to applied pressure. This enables the system to identify uneven pressure distribution by comparing the readings across the phantom.

The force-sensing resistors (FSRs) and Electrical impedance spectroscopy (EIS) were calibrated using a known standard calibration method. The FSRs were calibrated to ensure accurate pressure-to-resistance conversion. A custom-built calibration stand was used to apply a series of known weights ranging from 1 lb, 1.5lbs, 2 lbs, 5lbs (Jung et al., 2024).

The voltage output from the FSRs, which were configured in an inverted voltage divider circuit with a 10 kΩ reference resistor, was recorded using an ADS1115 analog-to-digital converter. The voltage readings were then used to calculate the corresponding resistance values. This data was used to generate a linear approximation model to convert subsequent resistance readings into quantifiable pressure units in kilopascal (kPa), ensuring the sensor’s output was accurate and repeatable. The EIS circuit was calibrated to validate its accuracy across the tested frequency range. This was a critical step because the circuit’s impedance measurement can be affected by external factors and hardware limitations.

The calibration was performed using a 10 kΩ reference resistor to represent a known constant impedance. The system was swept across its operational frequencies which were logarithmically spaced from 100.0 kHz, 122.3 kHz, 149.5 kHz, 182.9 kHz, 223.6 kHz, 273.4 kHz, 334.4 kHz, 408.9 kHz, and 500.0 kHz, Abasi et. al, (2022). The frequencies fall within the range of the beta dispersion region. In this region, the electrical current begins to penetrate cell membranes, allowing the EIS to detect differences in cell density and structure between healthy tissue and a simulated lump. The logarithmic sweep ensures a higher-resolution data collection at the lower frequencies, where the most significant impedance changes occur, without wasting time on the more gradual changes at higher frequencies.

### IV. Developing Simulated Breast Model and the Suspicious Breast Lump

The researcher developed a simulated breast model using 10% gelatin (50□g gelatin, 4.2□g agar powder, 1.25□g NaCl, and 361□mL water). The utilization of gelatine, agar, salt and graphite in simulated breast models aligns with the previous studies of Zengin and Gencer, (2025) these materials in replicating breast tissue properties and creating conductive inclusions for realistic lump simulation. In order to simulate a lump, a separate mixture of 3□g agar, 47□mL water, and 0.1□g graphite was prepared and embedded into the gel to mimic a suspicious breast lump for detection practice. The gelatin mixture without a lump was poured into a mold to serve as the control group (normal breast), while the mixture with the embedded lump served as the experimental group (breast with a suspicious lump), essential for testing the device.

### V. Testing Device on Simulated Breast

To evaluate the E-HAPLOS system, we conducted a series of tests on simulated breast phantoms to quantify its performance on classifying the lump. The experiment was performed using two gelatin-based phantoms: one representing healthy tissue and another with a simulated lump.

For the Force-sensing resistors, a total of 40 trials were performed to assess the relationship between the presence of the lump and the Force-sensing resistors. The researcher conducted 20 trials of each condition of the simulated breast with 2 different conditions: “With lump” and “Without lump.” (Boudewyn et al., 2017).

A total of 400 trials were conducted to assess the system’s ability to distinguish between the two tissue types (Boudewyn et al., 2017). This included 200 trials for the Electrical impedance spectroscopy (EIS) and 200 trials for the integrated EIS and Force-Sensing Resistor (FSR)-guided system.

Within each of these 200-trial sets, 100 trials were performed on the healthy phantom, while the remaining 100 trials were conducted on the phantom containing the simulated lump. This allowed us to evaluate the system’s performance under two distinct electrode placement conditions: on-center, where the EIS electrodes were guided to the location of the lump by the FSRs, and off-center, where the electrodes were placed without FSR guidance to simulate a non-optimized measurement.

### VI. Data Recording and Analysis

During the trials, the ESP32 microcontroller transmitted real-time pressure and impedance data to a laptop. This data was logged and stored as a .csv file for offline analysis.

The data analysis was performed using **Python** with the **pandas** and **scikit-learn** libraries. The mean and standard deviation of the impedance magnitude were calculated for each of the 100 trials per condition at each tested frequency.

The primary analytical methods included:

- **Descriptive Statistics**: The mean and standard deviation of both the impedance magnitude and the FSR pressure readings were calculated for each experimental condition (with and without a lump). 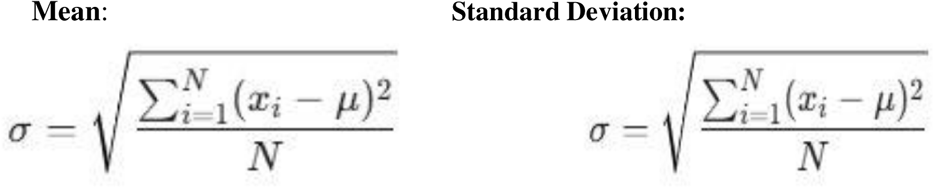
- **Logistic Regression and ROC-AUC Analysis**: Receiver Operating Characteristic (ROC) curves were generated using scikit-learn to evaluate the diagnostic performance of the EIS component. The Area under the curve (AUC) was calculated to provide a single metric for the system’s sensitivity and specificity. 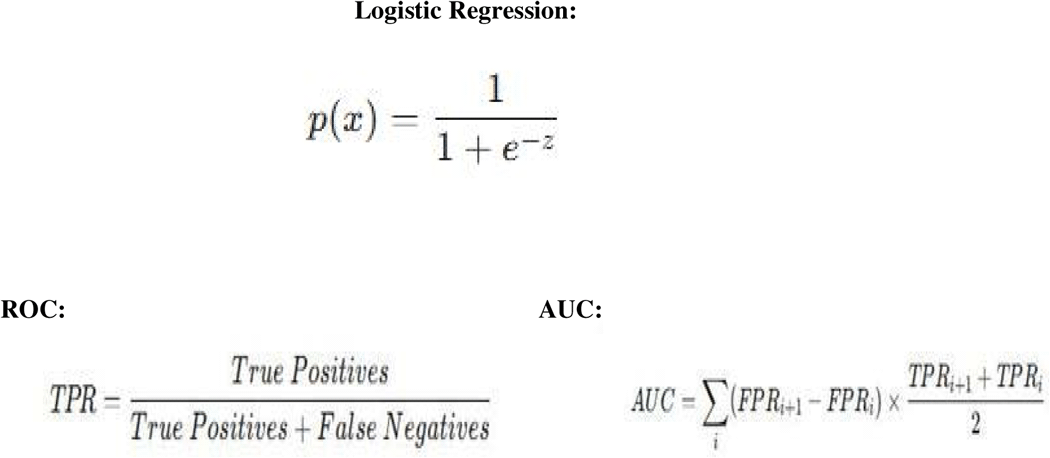
- **Two-Way ANOVA**: A Two-Way Analysis of Variance (ANOVA) is a statistical test used to determine how two independent factors, such as System Modality (FSR-guided, Unguided, or FSR only) and Frequency, affect a dependent variable, which in this case is the AUC score. This test is more powerful than a one-way ANOVA because it can analyze the main effects of each factor individually and, most importantly, the interaction effect. A significant interaction effect would be strong statistical evidence that the combined FSR-EIS system is uniquely effective. 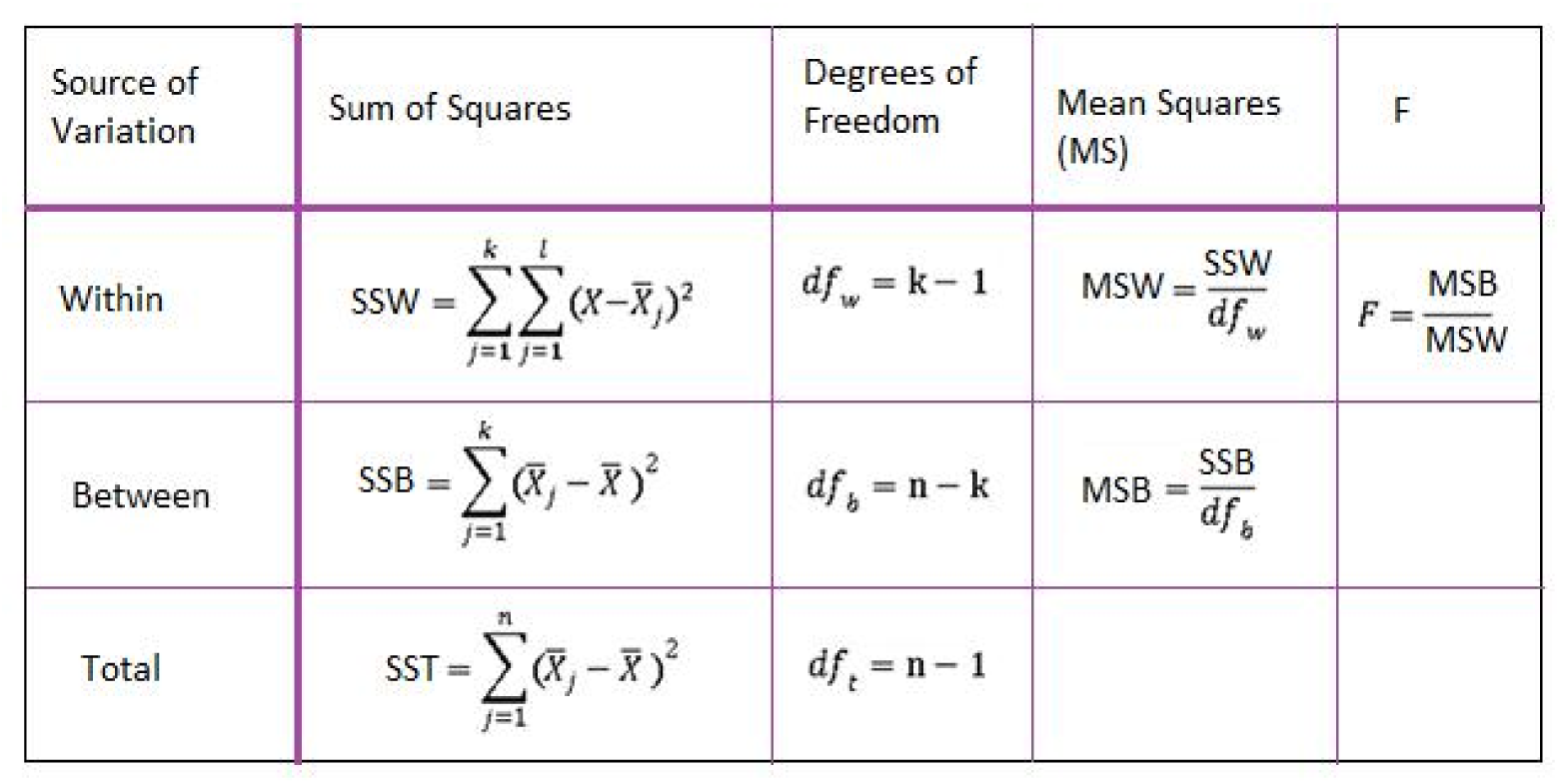
- **Post-Hoc Tukey’s HSD Test:** A post-hoc test, such as Tukey’s HSD (Honestly Significant Difference), is only used after a significant ANOVA result using the AUC score’s mean then subtract it to each other. Its sole purpose is to perform pairwise comparisons between your groups to identify exactly which groups have a statistically significant difference. It will confirm if your FSR-guided EIS system is statistically superior to your unguided EIS and FSR-only systems.

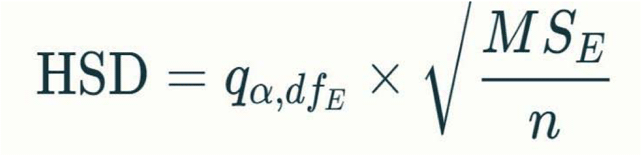

To validate the accuracy of the EIS circuit, a high-precision 10 kΩ reference resistor was used as a known constant impedance. The system was swept across its operational frequencies, from 100 kHz to 500 kHz, and the measurement error was calculated using the formula:

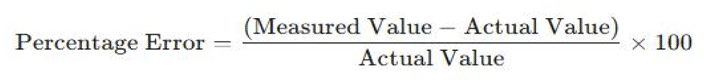

The calibration of the EIS circuit was a critical step to understand and quantify its accuracy across the operational frequency range. At the lowest tested frequency of 100kHz the circuit demonstrated high accuracy with a minimal error of approximately -2.5% indicating system’s reliability for measurement in this range. However, as the frequency increased, the measurement error grew substantially. By the time the circuit reached 500kHz, the error had increased to nearly -28%. This consistent negative error—where the measured impedance is lower than the actual value, it is primarily caused by parasitic capacitance. This is an unavoidable physical property in any circuit due to the close proximity of components and wires. At high frequencies, this parasitic capacitance creates a low-resistance path that allows the AC current to bypass the intended measurement components. This “short-circuiting” effect reduces the overall measured impedance, leading to the observed negative error.

The researcher acknowledged this as a limitation of the current design. The consistency of this error across all trials means that the system’s ability to discriminate between healthy and simulated lump tissue is not compromised, as the relative difference in impedance is still accurately measured.

### Risk and Safety

After the experiment, all electronic components such as soldering and wiring materials must be stored properly to avoid minor burns or injuries. The researcher must follow proper soldering techniques, dispose of wiring materials correctly, and handle sharp tools like scissors and wire cutters with care, as these pose a medium likelihood of cuts or punctures. Electrical hazards, such as short circuits and overheating, may cause minor shocks, burns, or sensor damage but can be prevented by insulating circuits, checking connections, using low-voltage power, and monitoring device temperature. Prolonged glove use may also cause skin irritation, which can be reduced by using breathable materials and allowing rest periods. According to Mohajan, et al. (2025) electronic components is one of the fastest and most growing waste streams in the world and it is classified as hazardous waste because of the expansion of the technological advancements, industrialization process, and higher living standards that poses a hazard to the ecosystem.

Maintaining safety protocols—such as wearing protective gear, organizing tools, and preventing overcrowded workspaces—protects the researcher and ensures reliable results. While risks are generally low, consistent caution and preventive practices are essential to safeguard both human safety and research integrity.

## RESULTS

### Test Results

**Table 1.**
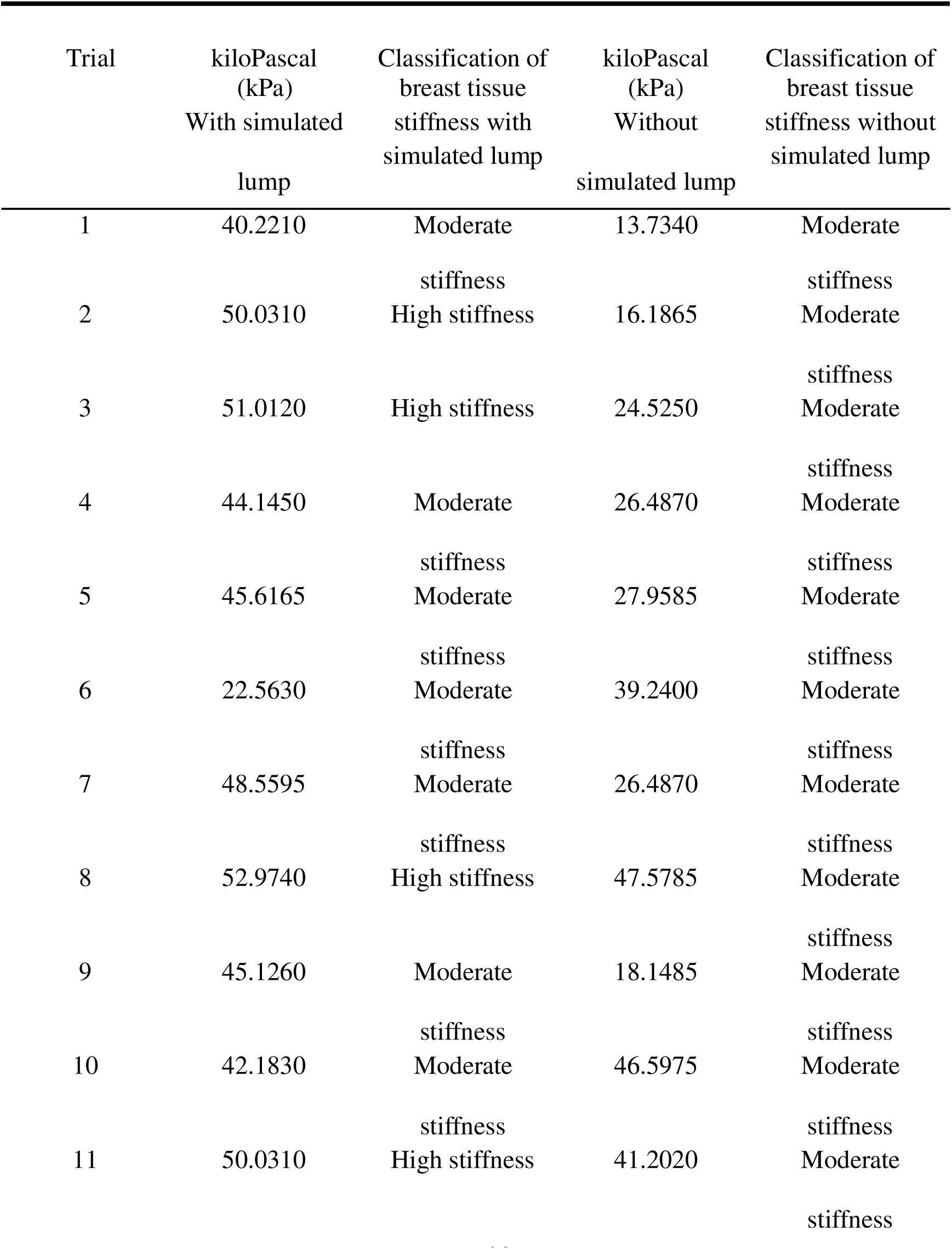
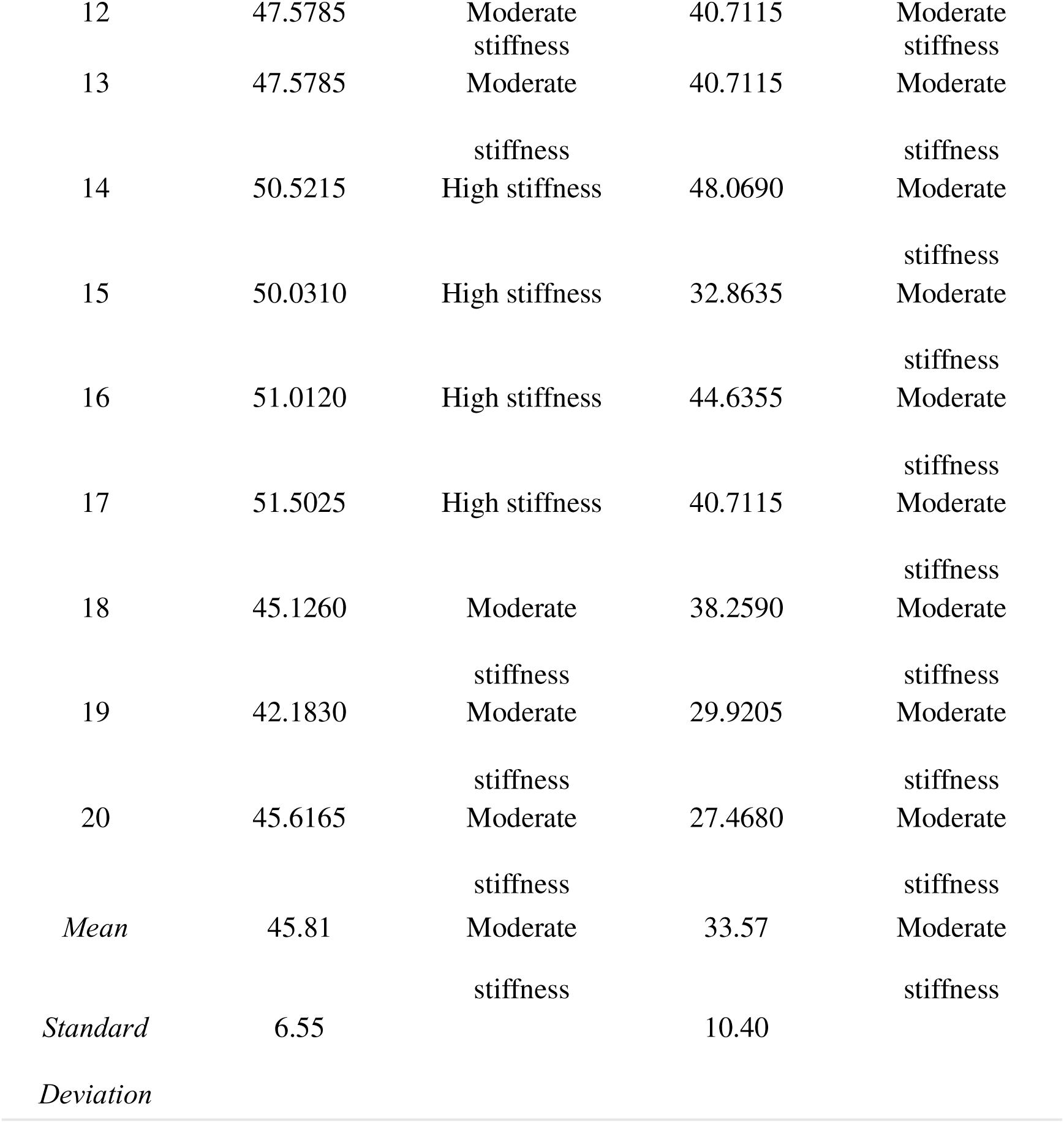
Force-sensor readings result from breast with and without simulated lump using logistic regression and visualizing using ROC curve and AUC scores.

**Figure 2.**
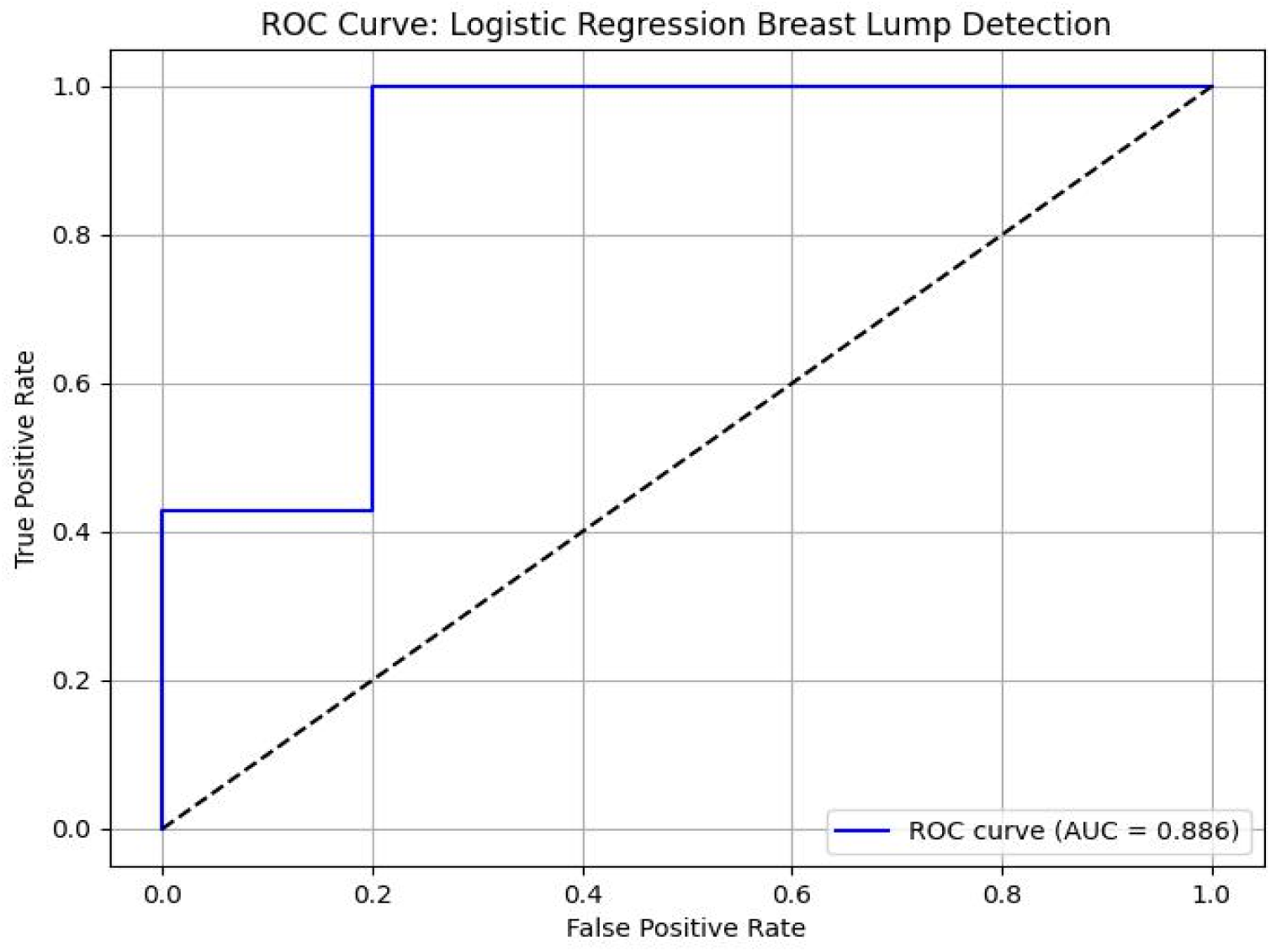
Logistic Regression Analysis: A Predictive Model for Stiffness in a unit of kilopascal (kPa).

**Table 2.** Summary of Measurement Error of EIS with 10kΩ Resistor.

| Frequency (Hz) | Measurement Error |
| --- | --- |
| 100,000 Hz | -2.53% |
| 122,280 Hz | -3.35% |
| 149,529 Hz | -4.57% |
| 182,875 Hz | -6.36% |
| 223,607 Hz | -8.64% |
| 273,397 Hz | -11.82% |
| 334,377 Hz | -16.08% |
| 408,890 Hz | -21.59% |
| 500,000 Hz | -27.92% |

## DISCUSSION

The findings confirm that stiffer tissues exhibit a significantly higher mean force as shown in Table 1, as evidenced by the measurements from the phantom with a simulated lump (Mean = 45.81 kPa, SD = 6.55) compared to the healthy phantom without lump (Mean = 33.57 kPa, SD = 10.40). This foundational result validates the use of Force-Sensing Resistors (FSRs) as an effective proxy for tissue stiffness. Figure 2 clearly indicates that the FSR correctly classifies the breast phantom, with ‘1’ representing the positive class (‘with a lump’) and ‘0’ representing the negative class (‘without a lump’).

The significant drop in AUC scores presented in Table 3 summarizing the score of ROC-AUC with a score ranging from 0.790 at 100kHz to 0.786 at 500kHz during the unguided EIS trials, which simulated an EIS measurement without knowing the exact location of the lump. The ROC curve was then visualized by Figure 3 showing the trend in the middle left indicating that the system struggles to identify the lump.

**Figure 3.**
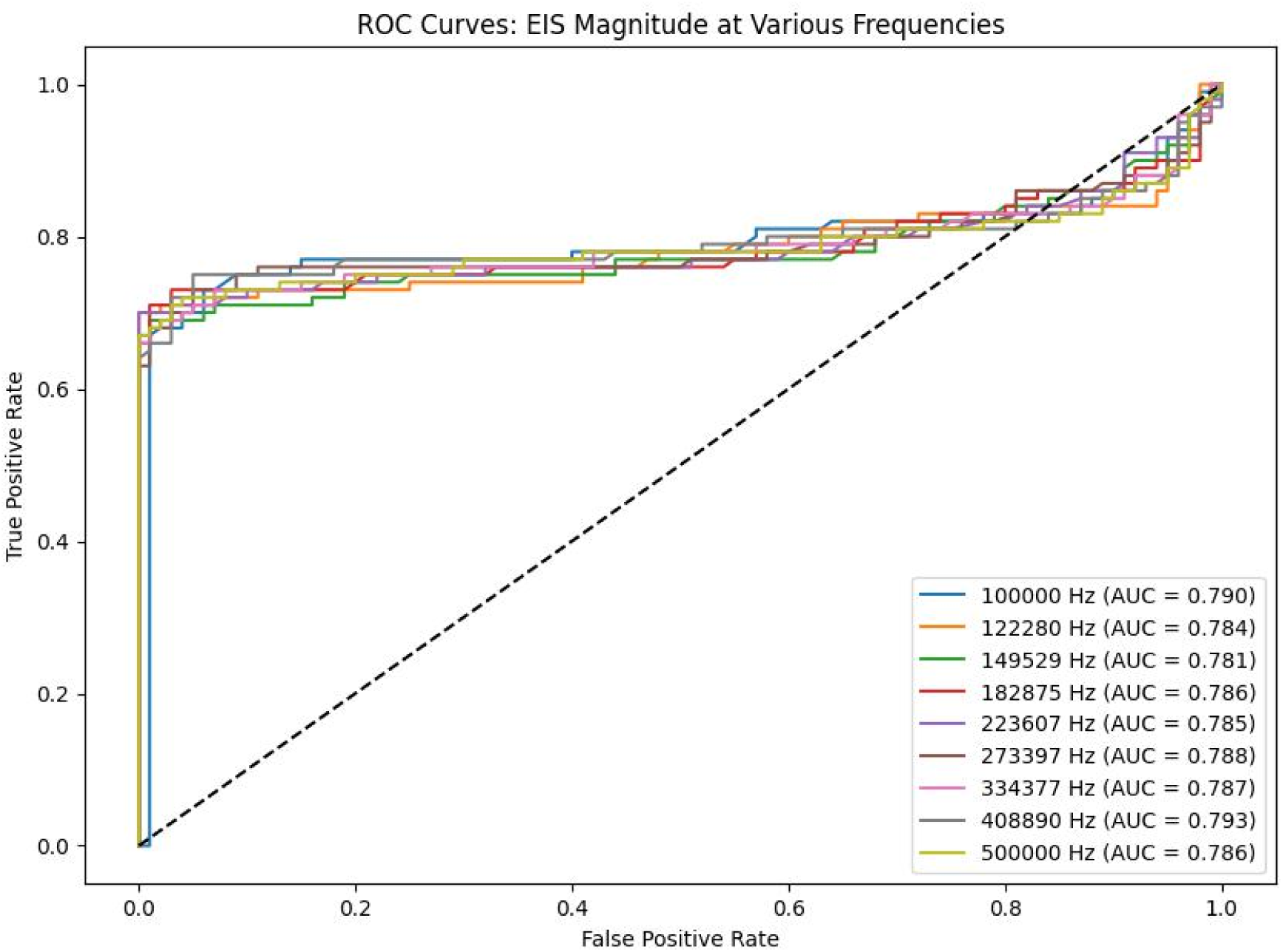
Receiver Operating Characteristic (ROC) curves for Unguided EIS off-center electrode placement at various frequencies.

**Table 3.**
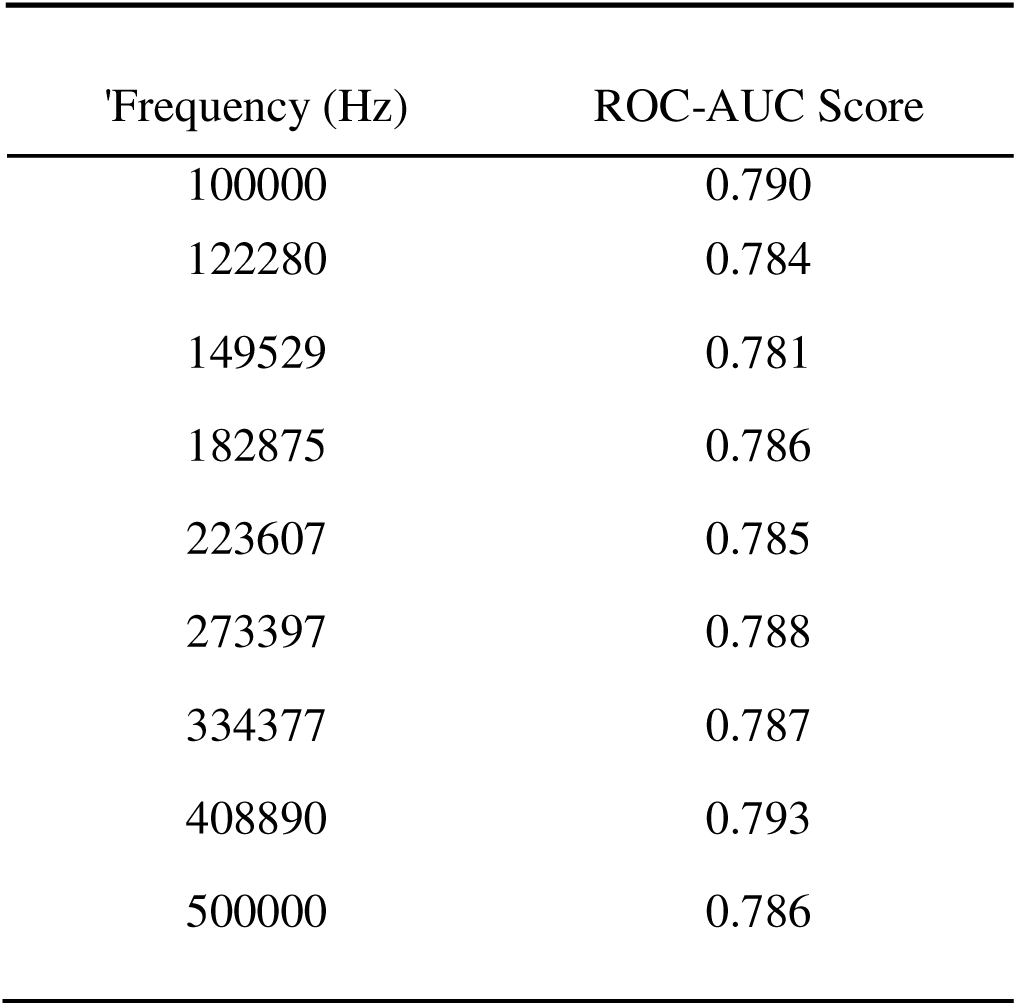
Summary of Unguided EIS Off-Centered ROC-AUC Scores.

The FSRs successfully guided the EIS electrodes to the optimal position in Figure 4, where the impedance contrast between the lump and surrounding tissue was most pronounced. This outcome supports the synergetic effect, as the combined system provides a significantly higher level of diagnostic performance, with AUC scores ranging from 0.944 to 0.951 as shown in Table 4 than the EIS modality without guidance.

**Figure 4.**
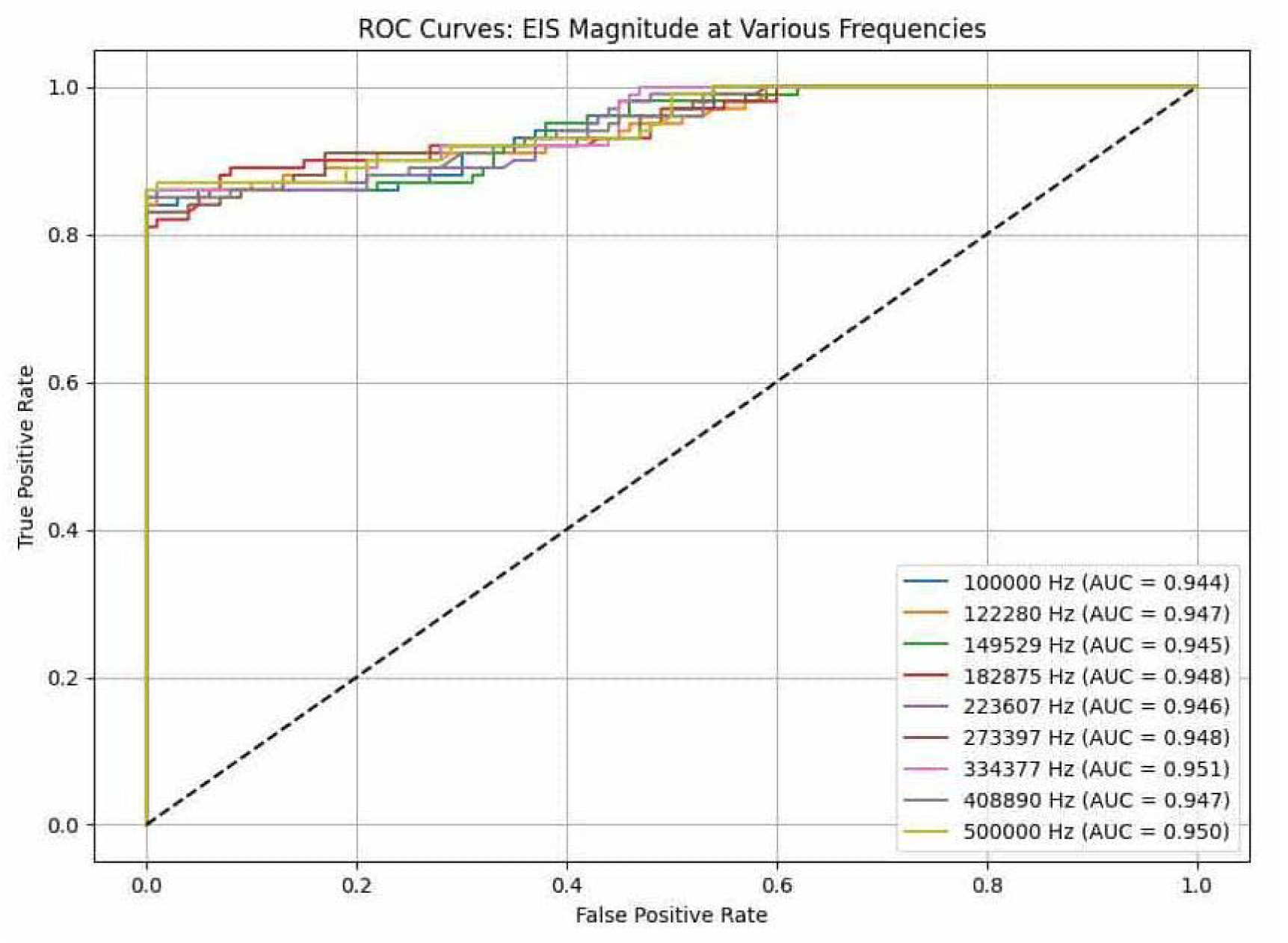
Receiver Operating Characteristic (ROC) curves for the FSR-guided EIS on-center

**Table 4.** Summary of Receiver Operating Characteristic (ROC) curves for the FSR-guided EIS on-center electrode placement.

| Frequency (Hz) | ROC AUC Score |
| --- | --- |
| 100000 | 0.944 |
| 122280 | 0.947 |
| 149529 | 0.945 |
| 182875 | 0.948 |
| 223607 | 0.946 |
| 273397 | 0.948 |
| 334377 | 0.951 |
| 408890 | 0.947 |
| 500000 | 0.950 |

The two-way ANOVA and Tukey’s HSD test provide a significant difference for this conclusion. The two-way ANOVA presented in Table 5 showed a significant difference between the diagnostic performances of the systems (p < 0.001), indicating a significant difference among the three groups. The FSR-EIS system had the highest performance, while the EIS-only system had the lowest. The Tukey’s HSD shown in table 6 confirmed that the FSR-guided EIS system was the primary driver of this difference with its performance being statistically superior to both the Unguided EIS and the FSR-only systems.

**Table 5.**
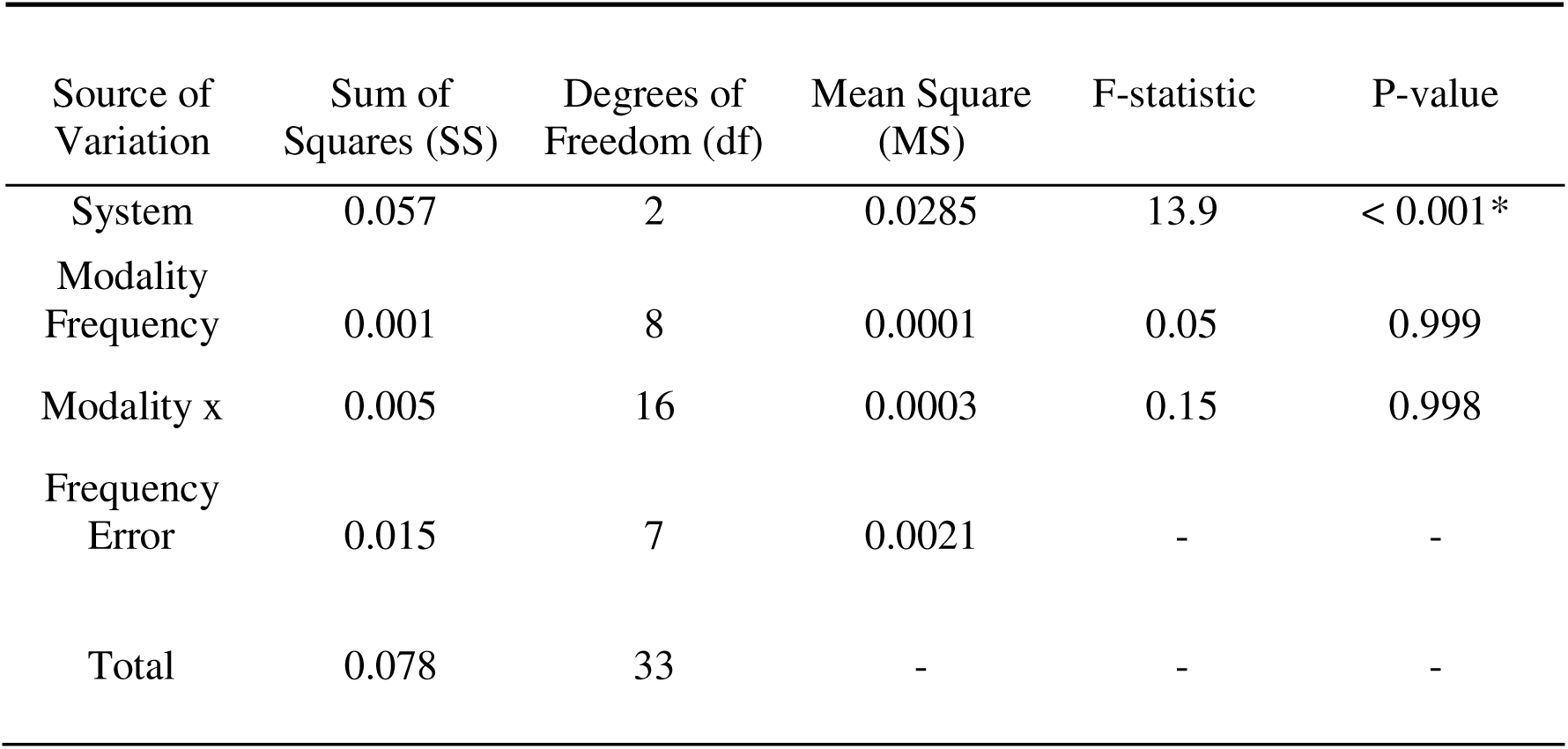
Two-Way ANOVA Results for Diagnostic Performance (AUC Scores)

**Table 6.** Post-Hoc Tukey’s HSD Test for Pairwise Comparisons.

| Comparison | Mean Difference | Standard Error | Tukey's HSD Q | P-value |
| --- | --- | --- | --- | --- |
| FSR-guided EIS vs. Unguided EIS | 0.16 | 0.021 | 7.61 | < 0.001* |
| FSR-guided EIS vs. FSR only | 0.061 | 0.021 | 2.90 | 0.041* |
| FSR only vs. Unguided EIS | 0.099 | 0.021 | 4.71 | <0.001* |

The results allow this study to confidently reject the null hypothesis (H□) and accept the alternative hypothesis (H□), which states that the combined FSR-EIS system provides a significant difference improvement in detecting the simulated lump than the individual sensor modalities.

This concept of synergy is well-supported in medical research study by Du et al. (2017) investigated the use of bioimpedance spectroscopy (BIS) to discriminate human breast carcinoma from benign tumors. While their study combined two electrical parameters (R0/R_∞_ and fc) and E-HAPLOS combines a physical measurement (FSR) with an electrical one (EIS), both projects demonstrate that combining distinct data types leads to a powerful synergistic effect that surpasses the capabilities of either modality alone. This shared principle confirms that our approach is consistent with established and successful methods in the medical field.

## LIMITATIONS ON RESEARCH DESIGN AND MATERIAL

A key limitation of this study is its use of simulated breast models instead of real human tissue. While these models effectively mimic the electrical and physical properties of breast tissue, they do not account for the complexities of the human body, such as varying tissue densities, blood flow, and individual patient anatomy. Additionally, the electronic circuitry demonstrated a frequency-dependent measurement error that increased at higher frequencies due to parasitic capacitance, indicating a need for a more advanced circuit design to ensure greater accuracy and repeatability.

## CONCLUSIONS

The E-HAPLOS system successfully demonstrated the synergistic effect of combining Force-Sensing Resistors (FSRs) with Electrical Impedance Spectroscopy (EIS) for detecting simulated breast lumps. The combined system achieved a diagnostic performance with an Area Under the Curve (AUC) above 0.94, which was statistically superior to unguided measurements (AUC ≈ 0.78) and FSR-only systems. This robust finding provides strong statistical evidence that the integrated E-HAPLOS glove holds great potential as an accessible screening tool.

The two-way ANOVA confirmed a statistically significant difference in diagnostic performance based on the system modality (*p* < 0.001). This was further supported by Tukey’s HSD post-hoc test, which showed that the FSR-guided EIS system was more evident than both the unguided EIS (*p* < 0.001) and the FSR-only system (*p* = 0.041). This finding provides strong statistical evidence that the integrated E-HAPLOS glove holds great potential as an accessible screening tool for breast lumps.

## RECOMMENDATIONS

Based on the promising results of this study, we recommend several steps for future research and development to transition the E-HAPLOS glove from a laboratory prototype to a clinically viable device. To minimize measurement error, advanced circuit design should be implemented, such as optimized Printed Circuit Board (PCB) layout, for more precise impedance measurements. Additionally, future research should explore machine learning algorithms to develop a predictive model from a larger dataset, which would allow for a more nuanced interpretation of sensor readings and potentially improve accuracy. The ultimate recommendation is to conduct clinical trials with human participants to validate the system’s effectiveness in a real-world setting, a crucial step in confirming its diagnostic performance, safety, and practicality. Finally, E-HAPLOS can potentially be used in health center as a reusable, accessible, and affordable screening tool that can be better alternative to manual palpation combined with adequate training for using the glove.

## APPENDICES

**Figure 5.**
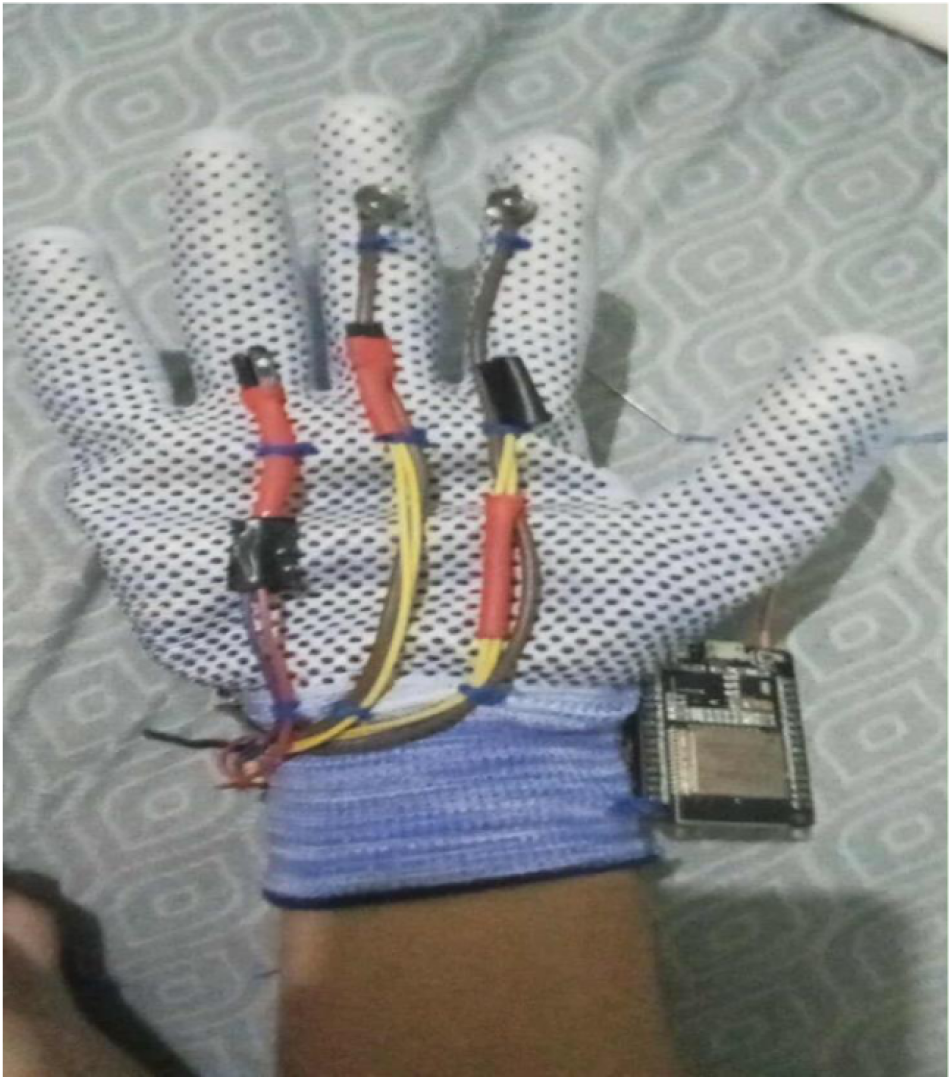
E-HAPLOS wearable gloves

**Figure 6.**
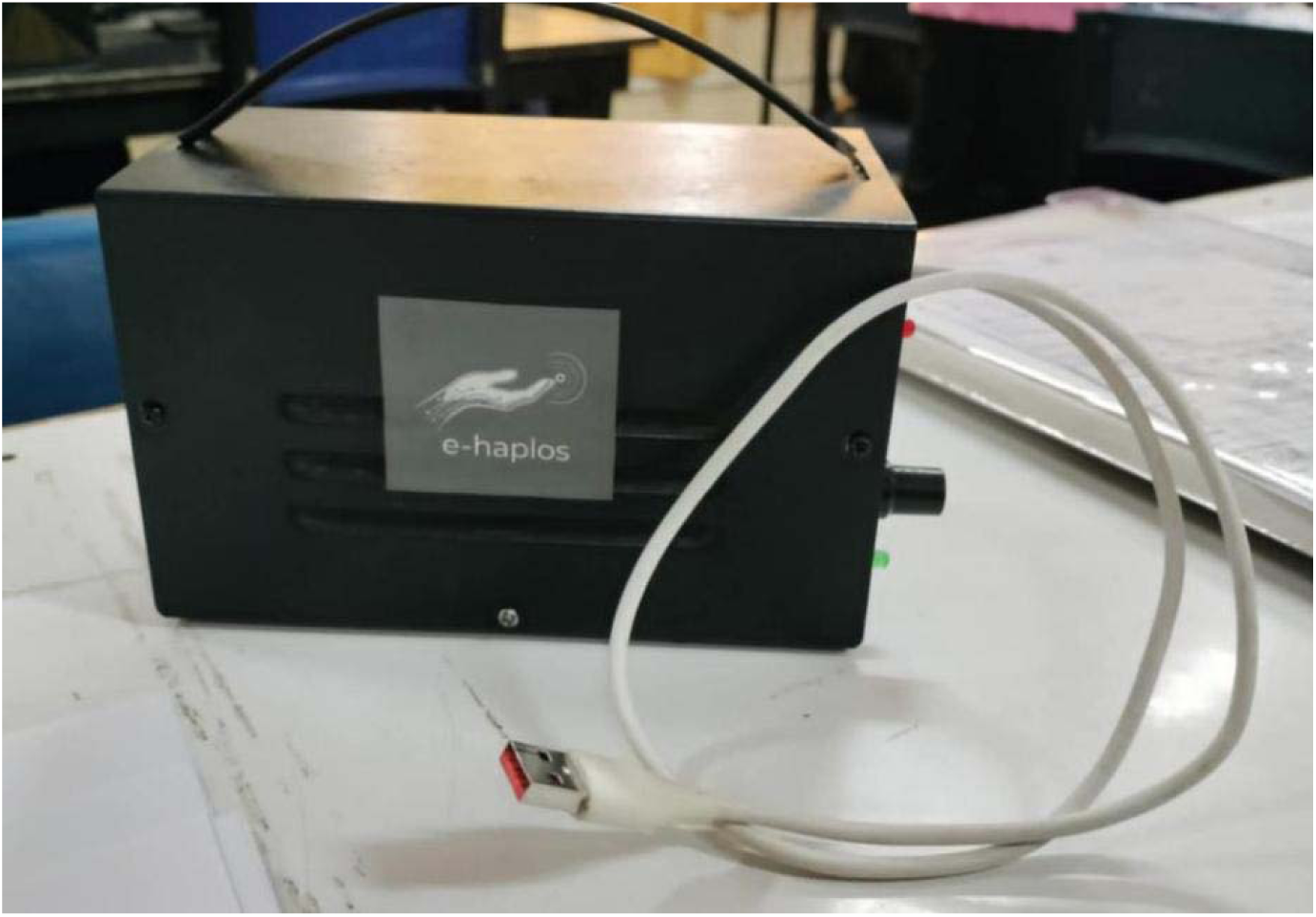
E-HAPLOS main processing unit

**Figure 7.**
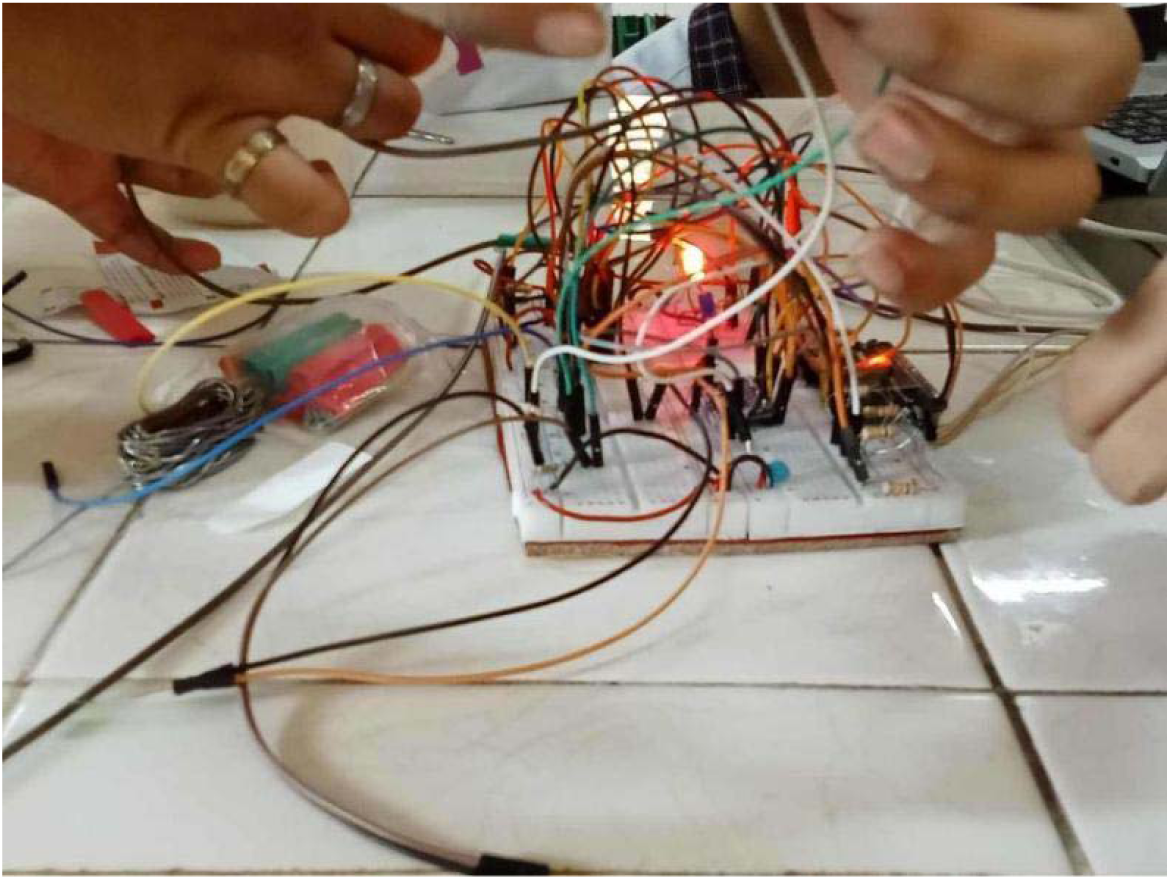
Calibrating the device

**Figure 8.**
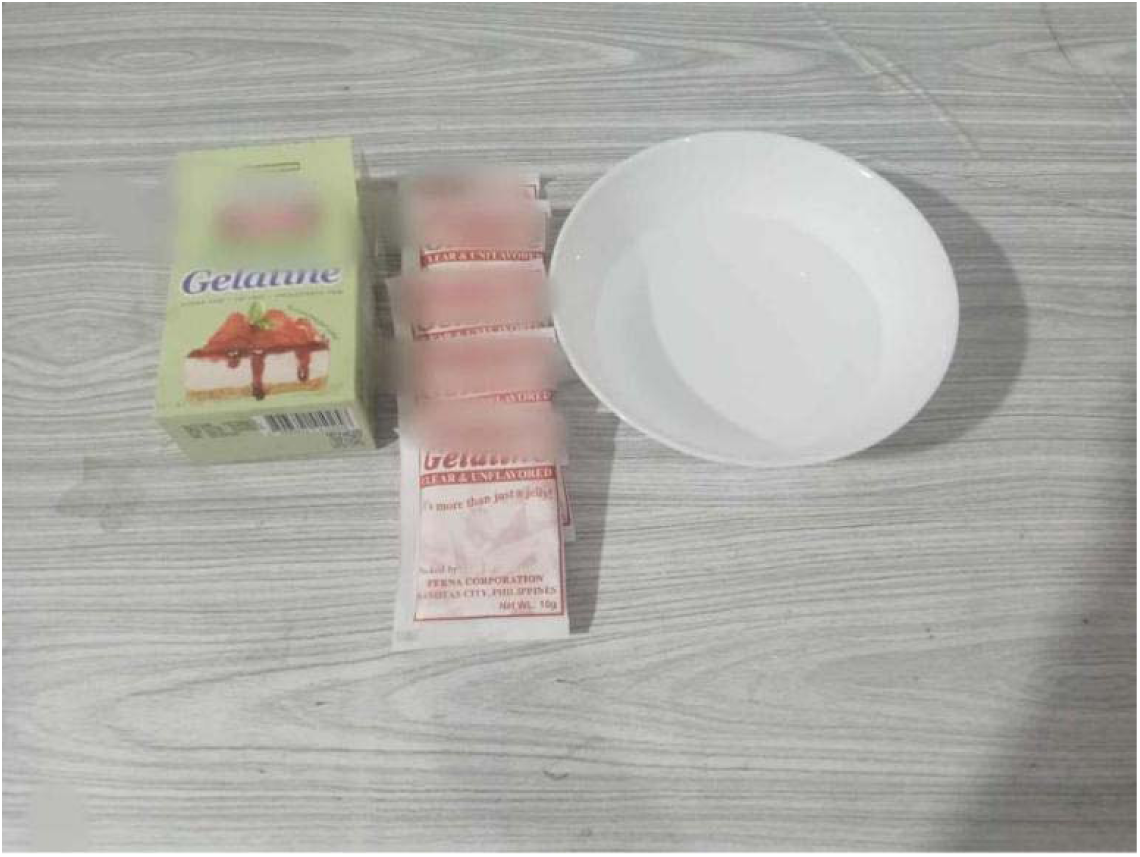
Components for making the breast model.

**Figure 9.**
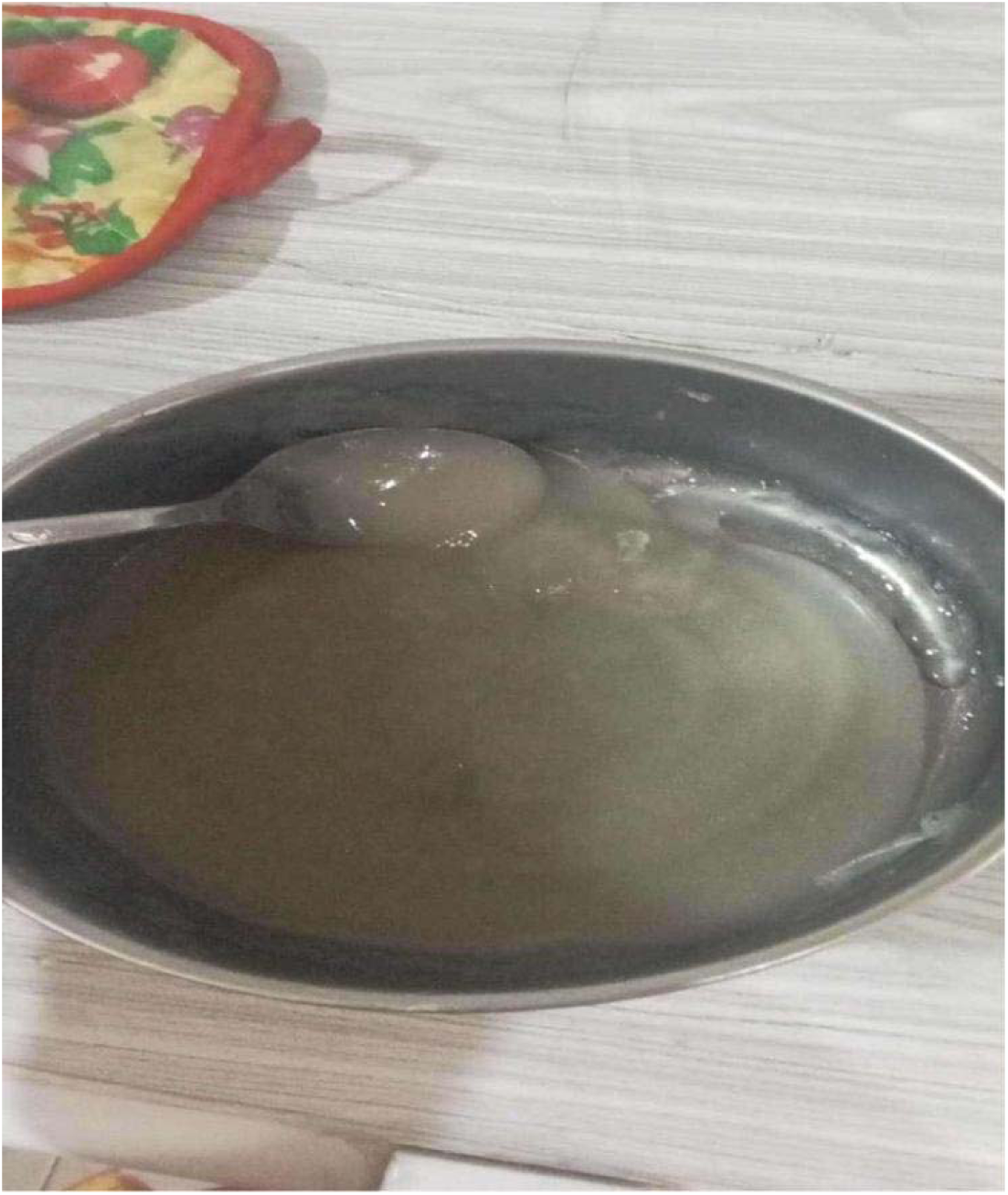
Mixing the gelatin to make a simulated breast model

**Figure 10.**
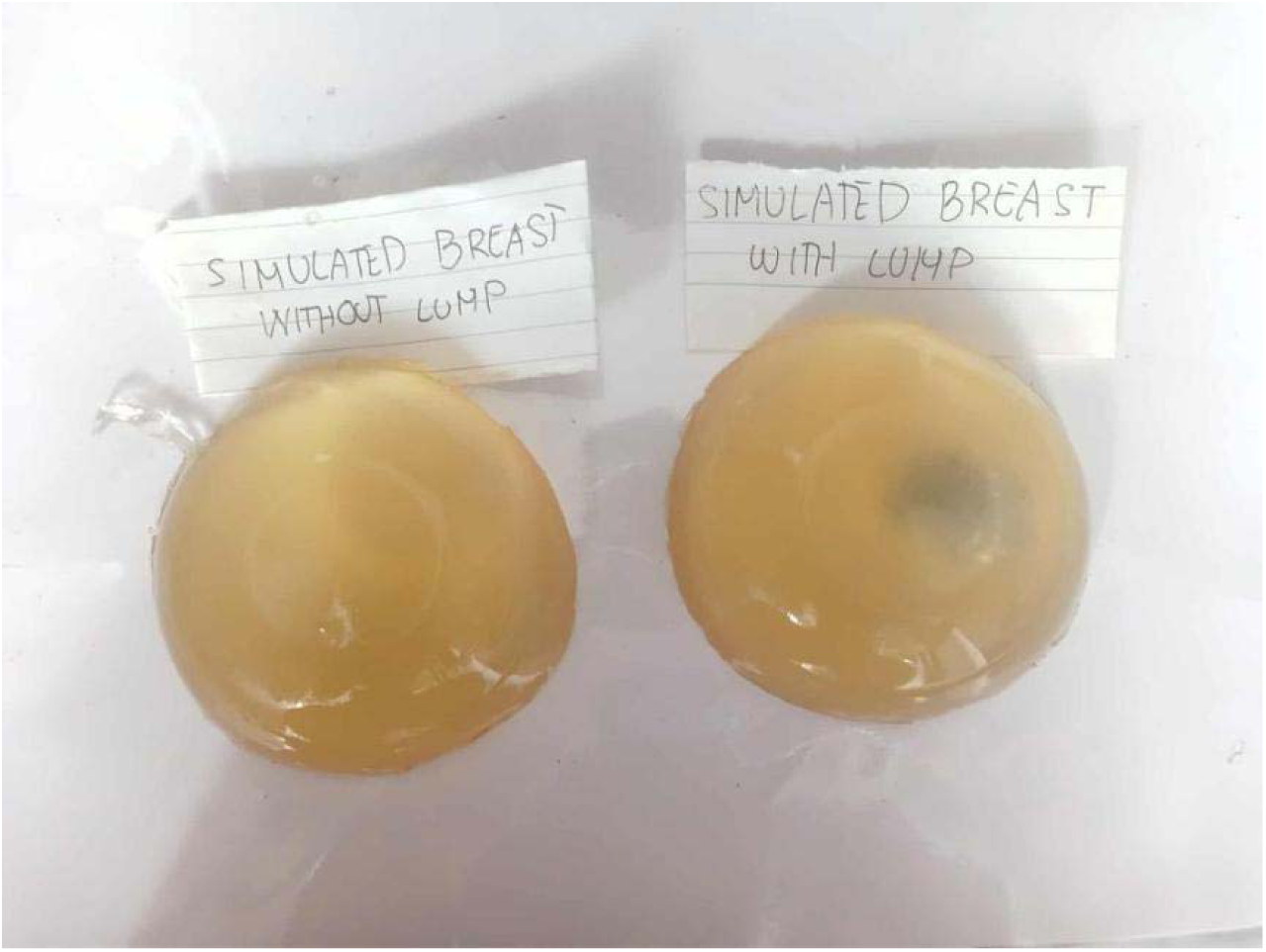
The simulated breast with lump and without lump.

**Figure 11.**
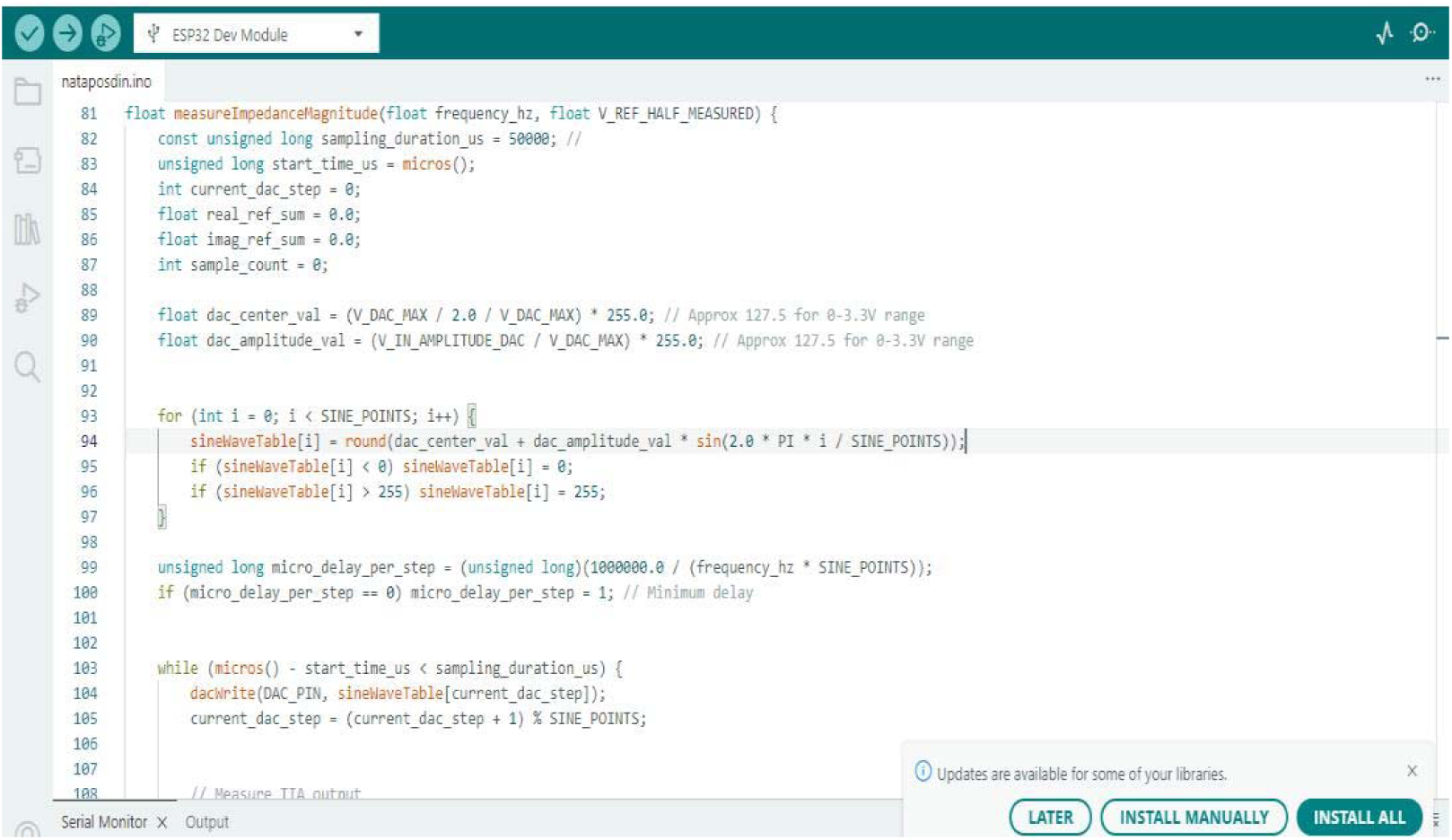
Codes for Electrical Impedance Spectroscopy C++

**Figure 12.**
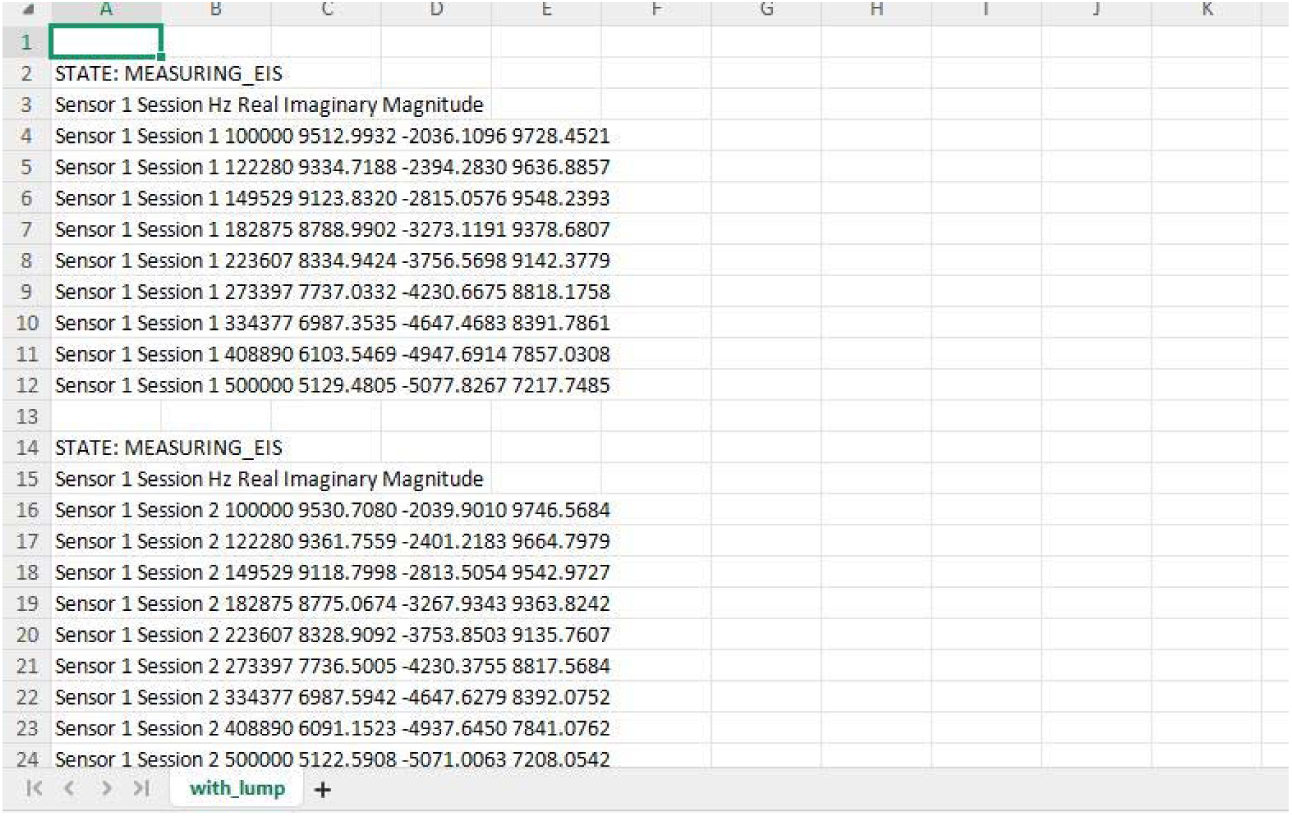
Readings of EIS during the experimentation of simulated breast with lump in .csv File.

